# Perceptual predictions track subjective, over objective, statistical structure

**DOI:** 10.1101/2025.07.30.667172

**Authors:** Jessye Clarke, Kirsten Rittershofer, Emma K. Ward, Daniel Yon, Clare Press

## Abstract

Over the past two decades, converging evidence from neuroscience and psychology has shown that predictions based on learnt statistical regularities exert a widespread influence on perception, action and cognition. Predictive processes in cognition and the brain are usually modelled as tracking objective event probabilities, deriving predictions and prediction errors from the statistical structure of the environment. However, our subjective models of our environments do not always align with these objective statistics. Currently we know little about how these subjective representations may determine the predictive functions. To separate subjective and objective contributions to prediction, we conducted three studies where cues (actions or tones) predicted visual outcomes (shapes or Gabors) with varying contingencies, and adult participants discriminated these outcomes. Uniquely to our paradigm, participants also reported their experiences of the statistical structure embedded in the task – the subjective probability (Experiment 1; *N* = 68), expectedness (Experiment 2; *N* = 35), or surprise (Experiment 3; *N* = 35) associated with the outcomes. When modelling subjective ratings alongside objective structure, the speed of perceptual decisions was best explained by independent, additive contributions of both. The decision itself was usually only explained by the subjective ratings, with little additional variance explained by objective statistical structure. These findings suggest that subjective experience may play a key, overlooked role in predictive processes, and open a host of interesting questions about the relative objective and subjective contributions to prediction, perception, and learning.

## 1. Introduction

The brain is often described as a ‘prediction machine’ (Friston, 2005, 2018), with human learners exquisitely sensitive to the statistics of their environment, and the resulting expectations shaping perception, action, and cognition in a multitude of interesting ways (Behrens et al., 2007; de Lange et al., 2018; Heilbron & Chait, 2018; Laukkonen & Slagter, 2021; Press et al., 2020; Seth, 2013). For example, statistical regularities are thought to generate predictions that bias us to perceive what we expect (Hudson et al., 2016; Wyart et al., 2012), with expected events reaching conscious awareness more rapidly (Pinto et al., 2015), and being discriminated more quickly and accurately (Stein & Peelen, 2015; see also Spaak et al., 2022). The influence of these regularities on cognition and neural processing is typically studied with respect to the objective statistics of the external environment, such as comparing high- and low-probability events (e.g., 75% vs. 25% as in Thomas et al., 2024; see also Garlichs & Blank, 2024) or modelling trial-by-trial changes in prediction error from normative models (den Ouden et al., 2009; Dijkstra et al., 2025). Relatedly, statistical learning is argued by some to be so ubiquitous that we are able to implicitly track the environmental statistics, even when we cannot report them (Turk-Browne et al., 2005, 2009). This line of work has been highly influential in explaining a range of cognitive functions.

However, these approaches largely neglect the fact that we often have reportable subjective experiences concerning the structure of our environment, and that these may play a role in the above predictive processes. We are certainly aware that red traffic lights are followed by yellow ones or that pressing a light switch will result in the room becoming illuminated. We also frequently experience a strong sense of surprise when events violate typical regularities – like when we intend to change gears on our bike and catch the wrong one or reach a sliding door that fails to open. Recent work indeed suggests that both implicit and explicit processes contribute to statistical learning, with distinct behavioural, neural, and memory outcomes depending on whether participants are aware of the regularities (Batterink et al., 2015; Conway, 2020). Interestingly, previous work has also suggested that subjective representations of statistical structure can depart from objective environmental statistics. For example, our probability judgements can differ systematically from the true probabilities of events, as exemplified by judging the conjunction of two events as more probable than the marginal probability of one of them (Tversky & Kahneman, 1983). Our reported surprise can also diverge from probability estimates, e.g., low-probability events appear unsurprising if they can be easily assimilated into existing models (Foster & Keane, 2015; Maguire & Maguire, 2009).

Appreciating the ways in which subjective representations may explain variance in predictive processes is thus crucial for understanding prediction, perception, and learning in their full complexity. It is currently unknown whether most predictive phenomena are modulated by objective statistical structure alone, or additionally or instead by subjective experience of it. Since one would assume that our subjective representations of probability and surprise are related to the objective statistical structure of our environments, the relative contributions of each are tricky to disentangle in current empirical work. In this work, we thus set out to test whether subjective experiences of statistical structure contribute to predictive influences on perceptual decision making.

We present three studies suggesting that perceptual predictions track subjective representations of statistical structure in ways not captured by the objective structure. Like prior work, we expose participants to statistical regularities between cues and visual stimuli, such that the stimuli vary in their expectedness based on the preceding cue. We ask participants to judge the identity of the stimuli to determine speed and accuracy of these perceptual decisions. Previous work using similar paradigms has shown that objective event probabilities reliably influence both reaction times and accuracy (Thomas et al., 2023). However crucially, in a twist on these usual paradigms, we also ask participants either at the end of the experiment to rate the probability of each cue-stimulus combination (Experiment 1), or to report on a trial-by-trial basis the extent to which they expected this stimulus (Experiment 2), or how surprised they were by its presentation (Experiment 3). We examined whether these subjective ratings explain variance in behaviour over and above that accounted for by the task’s objective statistical structure. To pre-empt our results, we find that decision speed is best explained with independent contributions from subjective ratings and objective structure, but decisions themselves are usually explained best solely according to the subjective ratings. These findings highlight the importance of considering subjective representations alongside objective statistics when investigating prediction, perception, and learning, and raise a host of interesting questions about their relative contribution to these functions.

## 2. Experiment 1

In Experiment 1, participants were exposed to statistical regularities between four cues (actions) and four different visual shapes. The conditional probability of a particular shape following a given action ranged from 71% to 2%. On each trial, participants were asked to judge the identity of the presented shape and at the end of the experiment, they rated the probability of each action-shape combination. Linear mixed effects models were fitted predicting reaction times and accuracy from either subjective probabilities, objective probabilities, or both.

### 2.1. Methods

#### 2.1.1. Participants

Sixty-eight participants (*M* age = 31.87 years, *SD* = 10.78) completed the study and were included in the analysis. An additional two participants did not meet our pre-registered inclusion criteria (overall mean reaction time above 1000 ms). All participants reported normal or corrected-to-normal vision and were recruited via Prolific. They gave informed consent prior to participation and were paid £3.50, equivalent to an hourly wage of approximately £7. The study complied with all relevant ethical regulations and was approved by the local Research Ethics Committee. The study design was pre-registered prior to data collection (https://doi.org/10.17605/OSF.IO/VRB2M).

#### 2.1.2. Procedure

The experiment was run online using Gorilla. On each trial, participants performed an action that triggered the presentation of one of four shapes. The objective probability of a particular shape following a given action varied across four levels (71%, 48%, 25% and 2%; see Fig. 1A), and participants had to judge its identity.

**Figure 1:**
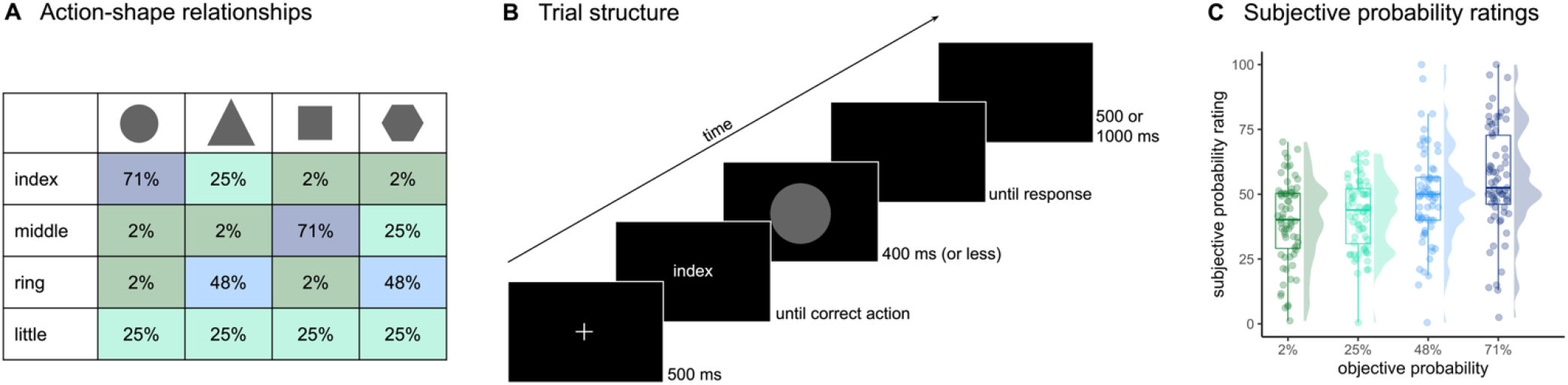
Experiment 1 design and subjective ratings. **(A)** Relationships between actions and shapes with the objective probability of a particular shape following a given action ranging from 71% to 48%, 25% and 2%. At the end of the experiment, participants rated the probability for each of these 16 action-shape combinations. **(B)** Trial structure. On each trial, participants were cued to perform an action and then presented with an outcome – one of the four shapes according to the relationships outlined in (A). Participants’ task was to indicate as quickly and accurately as possible which shape was shown. **(C)** Participants’ subjective probability ratings for the four levels of objective probability. As objective probability increases, so does rated probability. Coloured dots represent each participant’s mean rating at each level of objective probability. Boxes denote lower, middle and upper quartiles, and whiskers extend to 1.5 times the interquartile range. Half-violin plots display density estimates.

Each trial began with a fixation cross (500 ms), followed by the word ‘index’, ‘middle’, ‘ring’ or ‘little’, specifying which action to perform – a key press with the corresponding finger of their right hand. Once participants performed the correct action, a shape appeared on the screen and their task was to indicate as quickly and accurately as possible which shape was shown. The shape remained on the screen for 400 ms, or less if they responded before that. There was no time limit for responses, but participants were encouraged to respond before a warning tone played after 1000 ms. Participants responded using their left hand, pressing keys ‘1’ to ‘4’ for circle, triangle, square and hexagon, respectively. After responding, a blank inter-trial interval of either 500 or 1000 ms (pseudo-randomized) followed before the next trial began (see Fig. 1B).

Trial order within each block was randomized and participants received feedback on their accuracy and mean reaction time after each block. The experiment consisted of four blocks, each containing 100 trials. Each participant was pseudo-randomly assigned to one of twenty different randomly generated action-shape mappings.

After completing this task, participants were asked to rate the probability of each action-shape combination on a scale from ‘never (0%)’ to ‘always (100%)’ (e.g., ‘When you pressed down a key (‘H’) with your right index finger, how often was it followed by a circle?’). For all questions, the initial slider position was at the centre (50), and participants were required to move it away from and back to 50 if they wanted to select this value. Participants also rated how surprised they were by each action-shape combination on a scale from ‘0 (not surprised at all)’ to ‘100 (highly surprised)’, for which the results are reported in the supplementary materials. The same patterns emerged as with the subjective probability ratings.

#### 2.1.3. Analysis

The data were analysed using linear mixed effects models with the lme4 package in R (Bates et al., 2015), with all predictors being z-scored prior to analysis. Trials with reaction times (RTs) below 100 or above 2500 ms were excluded from all analyses. For RT analyses, only trials with correct responses were included.

To test whether participants’ subjective probability ratings were related to the objective probabilities, we fitted a linear mixed effects model predicting ratings from objective probabilities: lmer(subjective probability rating ∼ objective probability + (1 + objective probability | participant))

To test whether objective probabilities and subjective probability ratings explained participants’ RT and accuracy in responding to the shapes, we fitted three different models: a subjective model (subjective probability ratings as a fixed effect), an objective model (objective probabilities as a fixed effect), and a full model (including both measures as fixed effects). All models additionally included trial number as a fixed effect to account for global changes in behavioural performance throughout the experiment. For RTs, a linear mixed effects model predicted log-transformed RTs and for accuracy, a logistic mixed effects model predicted correct vs. incorrect response. See below for the full models:

lmer(log RT ∼ objective probability + subjective probability rating + trial number + (1 + objective probability + subjective probability rating + trial number ∣ participant)) glmer(accuracy ∼ objective probability + subjective probability rating + trial number + (1 + objective probability + subjective probability rating + trial number ∣ participant), family = binomial)

The subjective and objective models had the same random effects structure and only differed in their fixed effects.

We conducted model comparison using Bayesian Information Criterion (BIC) to assess which model – subjective, objective or full – provided the best overall fit to the behavioural data. Model comparison results using Akaike Information Criterion (AIC), which impose a less stringent penalty on complexity, are reported in the supplementary materials, but yielded similar conclusions to the BIC comparisons.

### 2.2. Results

Objective probability significantly predicted participants’ subjective probability ratings. The linear mixed effect model revealed a positive effect of objective probability on the ratings (*β* = 5.355, 95% CI [3.377, 7.333], *p* < .001), indicating that objectively more probable outcomes were also rated as more probable by the participants (see Fig. 1C).

To examine how both subjective and objective probability influenced participants’ behaviour, we fitted mixed effects models predicting RT and accuracy from either subjective ratings or objective probability. Crucially, we also included both predictors together in a full model to test whether each explained unique variance in behaviour. If both remained statistically significant, this would suggest that subjective ratings and objective probability independently contribute to behavioural responses.

When predicting reaction times, the subjective model revealed a negative effect of subjective probability ratings (*β* = -0.023, 95% CI [-0.034, -0.011], *p* < .001), indicating that participants responded faster for stimuli they rated as more probable at the end of the study. The objective model likewise showed a negative effect of objective probability (*β* = -0.024, 95% CI [-0.032, -0.016], *p* < .001), with faster RTs to objectively more probable outcomes. Importantly, the full model – containing both predictors – showed significant effects of both objective probability (*β* = -0.025, 95% CI [-0.033, -0.017], *p* < .001; Fig. 2A) and subjective probability ratings (*β* = -0.024, 95% CI [-0.036, -0.013], *p* < .001; Fig. 2C), suggesting that each explained independent variance in participants’ RTs. This pattern was also evident at the individual level: Most participants showed a negative slope between both objective and subjective probability and their RTs (see Fig. 2B and D). These results indicate that participants responded more slowly not only to outcomes that were objectively less probable, but also to those they judged as less probable. Model comparisons using BIC showed that the full model provided the best fit to the data (BIC = -9104.06), outperforming both the objective-only (BIC = -9099.10) and subjective-only (BIC = -9084.46) models.

**Figure 2:**
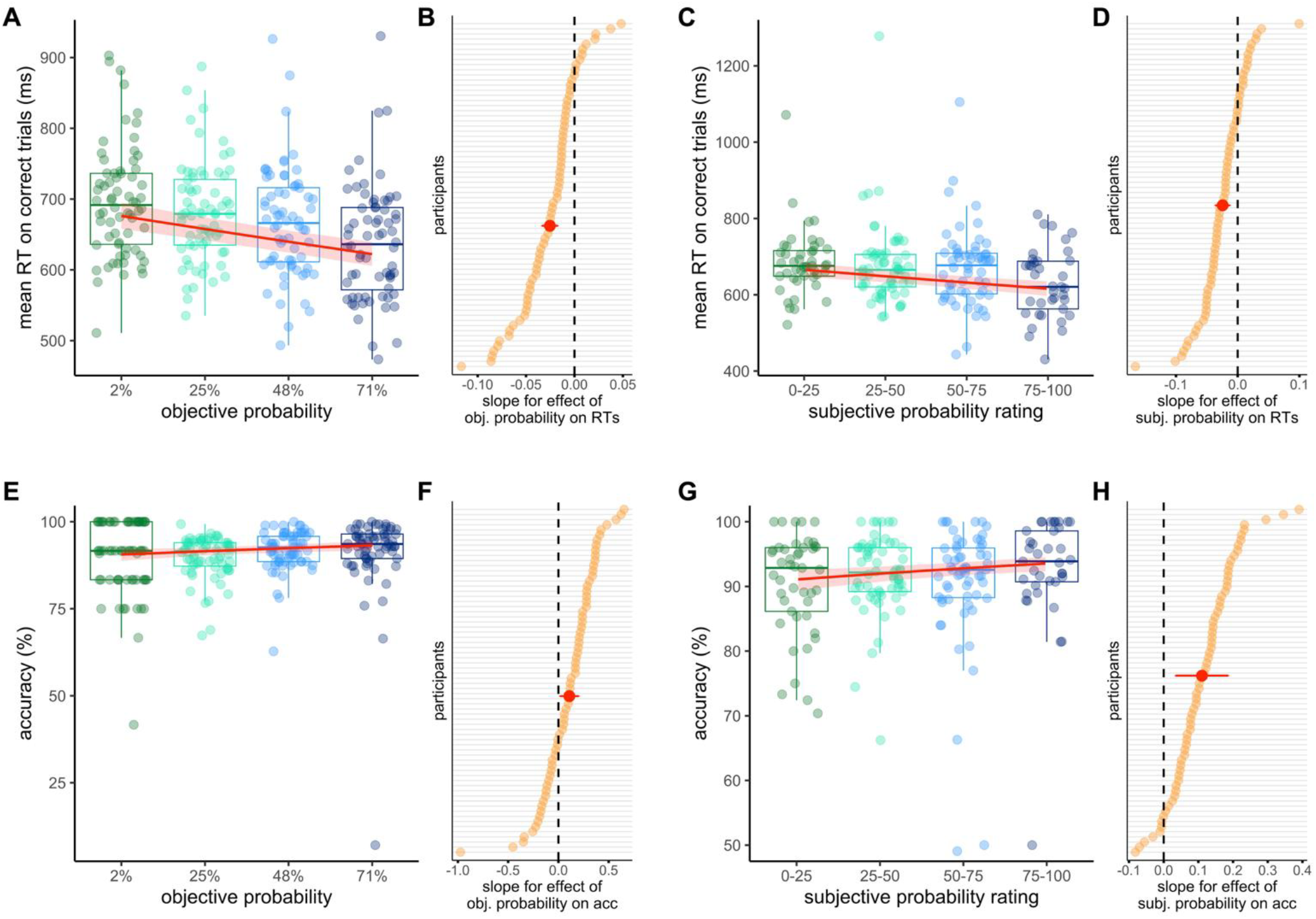
Experiment 1 results. **(A)** RTs as a function of objective probability, showing faster responses to outcomes that were objectively more probable. **(B)** Participant-specific slopes for the effect of objective probability on RTs, with most participants showing a negative slope. **(C)** RTs as a function of subjective probability ratings, indicating faster responses to outcomes rated as more probable. **(D)** Participant-specific slopes for the effect of subjective probability on RTs, with most participants exhibiting a negative relationship. **(E)** Accuracy as a function of objective probability, showing fewer errors for outcomes that were objectively more probable. **(F)** Participant-specific slopes for the effect of objective probability on accuracy, with the majority showing a positive association. **(G)** Accuracy as a function of subjective probability, showing fewer errors for outcomes rated as more probable. **(H)** Participant-specific slopes for the effect of subjective probability on accuracy, with the majority exhibiting a positive relationship. In (A), (C), (E) and (G), data points represent each participant’s mean performance within the four levels of objective probability or four equal-width bins of subjective probability ratings, created for visualization purposes only. Boxes show quartiles and whiskers extend to 1.5 times the interquartile range. Red lines represent the fixed effect estimates from the full model, with shaded ribbons showing 95% confidence intervals (CI). In (B), (D), (F) and (H), participant-specific slopes are displayed ordered by magnitude. Red points and horizontal bars indicate the fixed effect slopes and their 95% CI. Dashed vertical lines at zero indicate no effect.

When predicting accuracy, the subjective model revealed a positive effect of subjective probability ratings (*β* = 0.147, 95% CI [0.074, 0.221], *p* < .001), indicating that participants were more accurate for stimuli they judged as more probable. The objective model also showed a positive effect of objective probability (*β* = 0.158, 95% CI [0.066, 0.249], *p* < .001), suggesting higher accuracy for more objectively probable outcomes. In the full model, both subjective ratings (*β* = 0.110, 95% CI [0.035, 0.186], *p* = .004; Fig. 2G) and objective probability (*β* = 0.109, 95% CI [0.016, 0.201], *p* = .021; Fig. 2E) remained significant, with each explaining unique variance. This pattern was again evident at the individual level (see Fig. 2F and H). Model comparison this time however indicated that the subjective-only model provided the best fit (BIC = 15763.50), outperforming both the full model (BIC = 15768.56) and the objective-only model (BIC = 15765.89).

### 2.3 Discussion

These results suggest that participants’ reports of statistical structure explain unique variance in both the accuracy and reaction time of their perceptual decisions – beyond what is accounted for by the objective probabilities. In fact, when predicting accuracy, the model including *only* subjective rating and *not* objective probability was the best fit.

## 3. Experiment 2

Experiment 2 extended Experiment 1 by capturing how subjective experiences of statistical structure evolve during learning, employing trial-by-trial ratings rather than end-of-experiment judgements. We simplified the design such that participants were now exposed to regularities between just two cues and two outcomes, with cue-stimulus contingencies of 75% and 25%. We also chose a different subjective measure, such that participants rated now how much they had expected the outcome on each trial.

### 3.1. Methods

#### 3.1.1. Participants

Thirty-five participants (*M* age = 32.96 years, *SD* = 9.33) completed the study and were included in the analysis. An additional four participants did not meet our inclusion criteria – overall accuracy at the orientation discrimination task below 50%, or below 80% performance on the catch trials (see *3.1.2 Procedure*). All participants reported normal or corrected-to-normal vision, normal hearing, and were recruited from Prolific. They gave their informed consent prior to participation and were paid an hourly rate of £9. The study complied with all relevant ethical regulations and was approved by the local Research Ethics Committee.

#### 3.1.2. Procedure

The experiment was coded using JsPsych and run online using the MindProbe server. On each trial, participants were presented with one of two tones (330 Hz, 784 Hz) followed by either a clockwise (+45 degrees from vertical) or counterclockwise (-45 degrees) oriented Gabor patch. Action cues (Experiment 1) were replaced by tones to reduce the number of button presses on each trial, and shape stimuli were replaced by Gabor patches to render them of identical complexity. Each tone predicted one orientation with 75% probability and the other with 25% probability (see Fig. 3A). Participants had to judge the presented orientation as well as rate how much they had expected this orientation to be shown.

**Figure 3:**
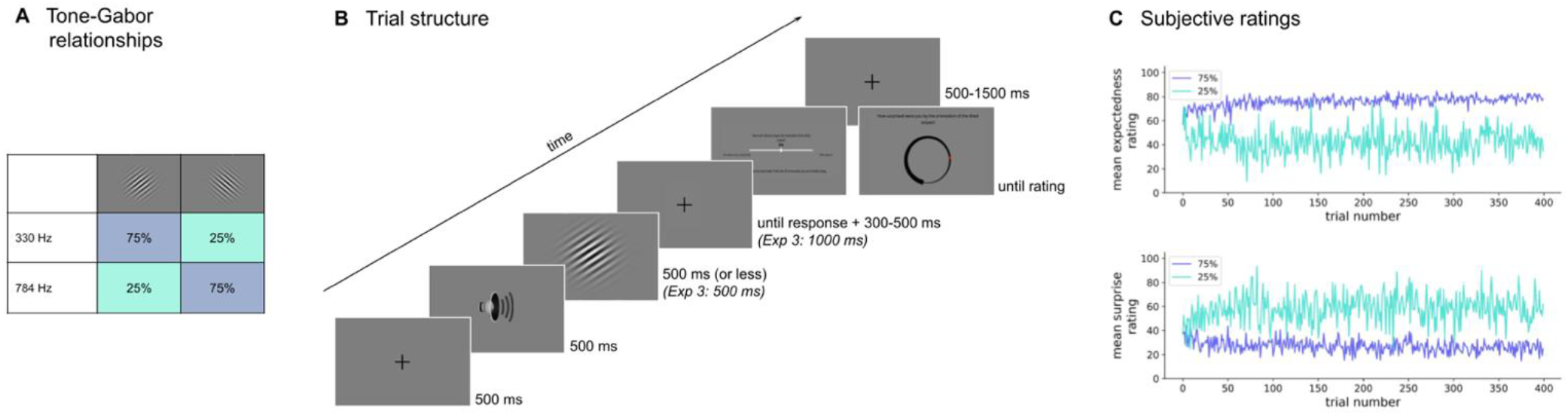
Experiments 2 and 3 design and subjective ratings. **(A)** Example mapping between tones and orientations. Each tone predicted one orientation with 75% and the other with 25% probability. **(B)** Trial structure. On each trial, participants were presented with a tone followed by a Gabor. They had to discriminate the Gabor orientation and also rated how expected (Experiment 2) or surprising (Experiment 3) they found the presented orientation. **(C)** Participants’ subjective expectation (Experiment 2) and surprise (Experiment 3) ratings over the course of the experiment for the 75% and 25% trials.

Each trial began with a fixation cross (500 ms), followed by a tone (500 ms) and the subsequent presentation of a Gabor. Participants’ task was to discriminate its orientation as quickly and accurately as possible using the ‘A’ and ‘Q’ keys. The Gabor remained on the screen for 500 ms, or less if they responded before that. There was no time limit for responding. After participants’ response, there was a pseudo-randomised blank screen of 300-500 ms. Next, they rated how much they had expected the presented orientation, using a continuous scale from ‘0 (very unexpected)’ to ‘100 (100% expected)’. The initial slider position was again at the centre (50), and participants had to move the slider at least once to continue. Each trial ended with a 500-1500 ms pseudo-randomised inter-trial interval (see Fig. 3B).

The tone-orientation mappings and the key-orientation responses were counterbalanced across participants. The experiment had 400 trials, and every 100 trials participants were given feedback on their accuracy, as well as a recap of the instructions. There were 20 catch trials, in which participants were asked whether two consecutive tones were the same or different, distributed randomly throughout the experiment to check that participants’ computer audio was working correctly.

#### 3.1.3. Modelling

To capture the objective statistical structure of the task on a trial-by-trial basis, we fitted a Rescorla-Wagner (RW) learning model (Rescorla & Wagner, 1972) to each participant’s trial sequence of cue-stimulus pairings. The model estimated the associative strength between the auditory cue and presented Gabor orientation on each trial, and these trial-wise values served as our estimates of objective expectations. The Rescorla-Wagner model is a widely used model of associative learning, and Rescorla-Wagner parameters have been shown to describe learning behaviour (Olawole-Scott & Yon, 2023; Roesch et al., 2012; Williams et al., 2017) and be represented across different neural structures (den Ouden et al., 2009; Rodriguez et al., 2006; Roesch et al., 2012). The model maintains separate expectation values (*V*) for each tone (*T*), initialized at 0 and updated on each trial using the following equation:

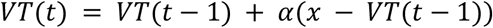

*VT*(*t*) represents the model’s expectation for the given tone *T* at trial *t*, and *x* is the presented orientation, coded as 1 (clockwise) or -1 (counterclockwise). Expectations are updated in response to presented stimuli at a magnitude controlled by the learning rate *α*. The learning rate was determined by minimizing the mean squared prediction error across each participant’s unique trial sequence. To achieve this, we tested 1,000 candidate α values between 0 and 1 and selected the one that resulted in the smallest mean squared prediction error. Participants’ behaviour is not used in this procedure: the best-fitting parameter values are determined solely by minimising the mean squared prediction error for the actual sequence of presented stimuli, independently of how participants responded.

When a tone is presented, the model makes a prediction about the expected orientation based on its current *VT* value. If the presented orientation matches the expected one, the model strengthens its expectation. If the presented orientation is unexpected, the model adjusts its expectation accordingly. For example, if Tone 1 is frequently followed by a clockwise (1) Gabor patch, over time, *VT*1 will move towards 1, meaning the model increasingly expects clockwise orientations when Tone 1 is played. Conversely, if Tone 2 is frequently followed by a counterclockwise (-1) Gabor patch, *VT*2 will move towards -1. On each trial, the model computes a prediction error as the difference between the presented orientation and the expected value:

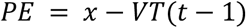

This prediction error determines the size of the expectation update, with larger errors leading to bigger adjustments. To derive a measure of expectation strength independent of stimulus orientation, we multiplied counterclockwise expectations by -1, ensuring that expectation values ranged from 0 to 1 for all trials. However, since these values reflected the expected stimulus given the tone – and an unexpected stimulus was shown on 25% of trials – we adjusted for these cases by computing (*VT* ∗ −1) on unexpected trials. This transformation resulted in a scale from -1 (highly unexpected) to 1 (highly expected), reflecting the expectedness of the presented stimulus rather than the expected stimulus. For example, if the model had an expectation of 0.8 for clockwise, but a counterclockwise stimulus was presented, the final expectedness value for this trial would be -0.8, indicating that the presented stimulus was unexpected. This provided a trial-by-trial measure of the objective expectedness of the presented orientation, given the tone.

#### 3.1.3. Analysis

For all analyses, RTs below 100 and above 2500 ms were excluded. For the RT analyses, trials in which the orientation discrimination question was answered incorrectly were additionally excluded.

As in Experiment 1, participants’ subjective trial-by-trial ratings were predicted from the objective model-derived expectedness values using a linear mixed effects model. For analysing participants’ behavioural performance, three different models were used to predict RTs and accuracy in the orientation discrimination task: a subjective model with participants’ trial-by-trial expectedness ratings, an objective model using the trial-by-trial objective expectedness values derived from the RW model, and a full model including both predictors. In the accuracy models, the inclusion of trial number as a fixed or random effect resulted in convergence issues. As trial number was not central to our theoretical predictions, we opted to remove it from these models to ensure model stability. Trial number was retained as a fixed and random effect in all reaction time models, where convergence was achieved. Model comparisons were again conducted using BIC, with AIC results reported in the supplementary materials and yielding similar conclusions.

### 3.2. Results

To test whether our design successfully induced learning and the development of perceptual expectations in participants, we first compared RT and accuracy scores for the orientation discrimination task in the 75% and 25% conditions. A paired t-test confirmed that responses were significantly faster (*t*(34) = - 6.274, *p* < .001) in the 75% (*M* = 790.55 ms, *SD* = 299.52) than in the 25% condition (*M* = 926.31 ms, *SD* = 330.23). Analysis of accuracy also confirmed it to be higher (*t*(34) = 3.171, *p* = .003) for 75% trials (*M* = 94.74%, *SD* = 8.72) than 25% trials (*M* = 91.18%, *SD* = 8.02). These analyses suggest that participants learnt the mappings between audio cues and Gabors.

Investigating the relationship between subjective expectedness ratings and Rescorla-Wagner derived expectedness/associative strengths, revealed a significant positive effect of RW expectedness on subjective ratings (*β* = 15.066, 95% CI [10.836, 19.296], *p* < .001), indicating that participants’ subjective expectedness ratings tracked the associative strength predicted by the Rescorla–Wagner model. Mean ratings for each trial are visualised in Figure 3C.

In the reaction time results, the subjective RT model revealed a negative effect of subjective expectedness (*β* = -0.134, 95% CI [-0.160, -0.108], *p* < .001), indicating that participants discriminated the orientation faster when they rated the stimulus as more expected. The objective model showed a negative effect of objective RW expectedness values (*β* = -0.062, 95% CI [-0.079, -0.046], *p* < .001), showing that RTs decreased as objective associative strength increased. The full model showed effects of both objective expectedness (*β* = -0.040, 95% CI [-0.057, -0.024], *p* < .001; Fig. 4A and B) and subjective expectedness (*β* = -0.109, 95% CI [-0.136, -0.081], *p* < .001; Fig. 4C and D), demonstrating that the factors of subjective and model-generated expectations explained independent variance. Model comparison using BIC indicated that the full model – including both subjective and model-derived expectedness – provided the best fit to the data (BIC = 8128.76), outperforming the subjective-only model (BIC = 8137.59) and the objective-only model (BIC = 8152.68).

**Figure 4:**
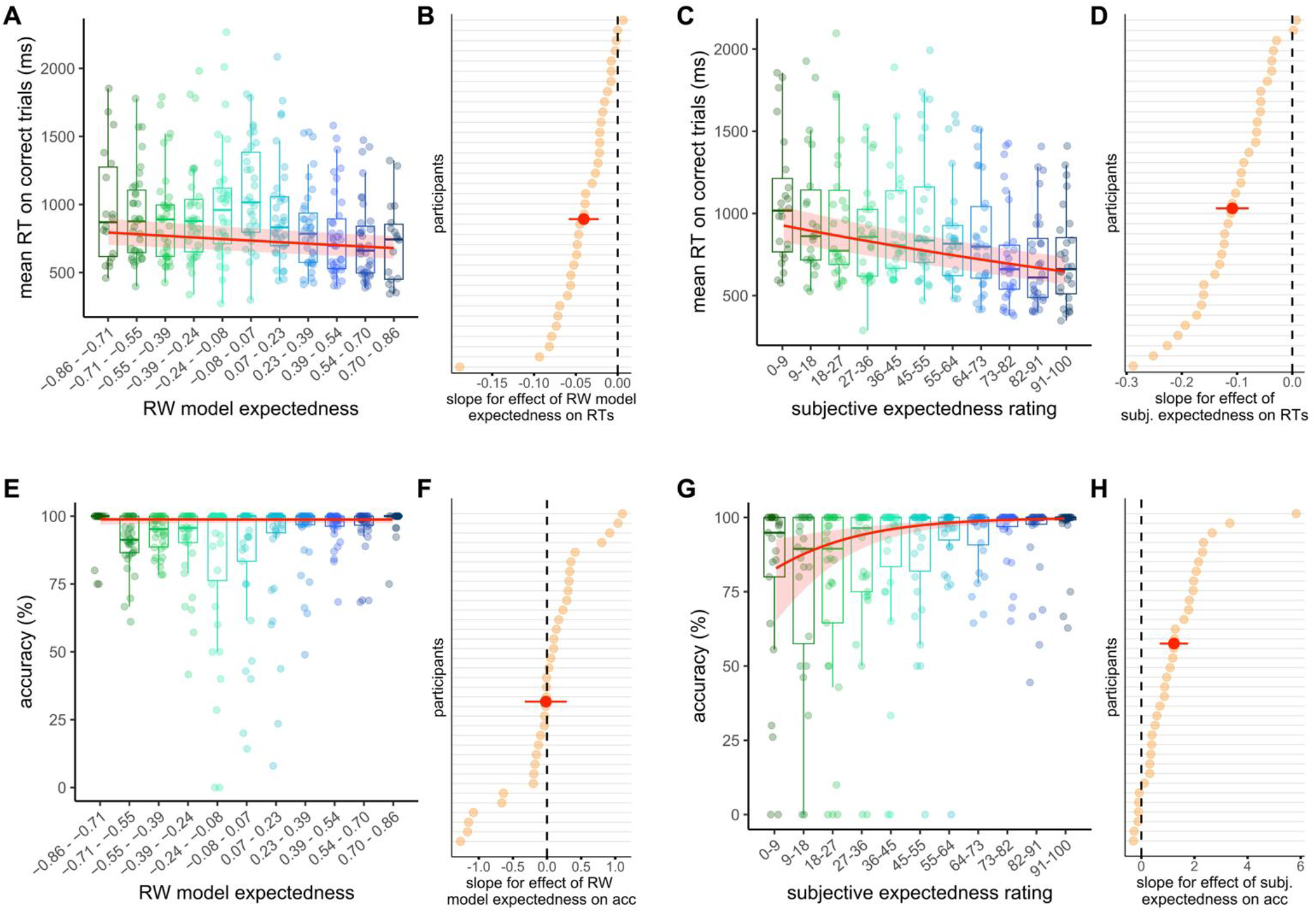
Experiment 2 results. **(A)** RTs as a function of Rescorla-Wagner expectedness, showing faster responses when there was a higher objective associative strength between outcome and cue. **(B)** Participant-specific slopes for the effect of RW expectedness on RTs, with most participants showing a negative slope. **(C)** RTs as a function of subjective expectedness ratings, indicating faster responses to outcomes rated as more expected. **(D)** Participant-specific slopes for the effect of subjective expectedness on RTs, with most participants exhibiting a negative relationship. **(E)** Accuracy as a function of RW expectedness. When included alongside subjective expectedness ratings, objective RW expectedness had no significant effect on accuracy. **(F)** Participant-specific slopes for the effect of objective RW expectedness on accuracy. **(G)** Accuracy as a function of subjective expectedness, showing increasing accuracy as outcomes were rated as more expected. **(H)** Participant-specific slopes for the effect of subjective expectedness on accuracy, with the majority exhibiting a positive relationship. Plotting conventions are the same as for Fig. 2. For visualization purposes, data was grouped into eleven equal width bins of either RW expectedness value (A, E) or subjective expectedness rating (C, G) and participants’ mean performance was calculated. Bins span the minimum to maximum observed values (RW expectedness can theoretically range from –1 to 1).

In the accuracy models, the subjective model showed that subjective expectedness rating was positively associated with accuracy (*β* = 1.211, 95% CI [0.801, 1.621], *p* < .001), indicating that as participants increasingly reported expecting the stimulus, performance at the discrimination task improved. Objective expectedness also predicted accuracy (*β* = 0.402, 95% CI [0.089, 0.715], *p* = .012), suggesting that participants were more likely to respond correctly as model-generated expectedness increased.

However, in the full model, while subjective expectedness significantly predicted accuracy (*β* = 1.228, 95% CI [0.731, 1.724], *p* < .001; Fig. 4G and H), objective expectedness did not (*β* = -0.017, 95% CI [-0.309, 0.275], *p* = .907; Fig. 4E and F), suggesting that in this case, the variance explained by objective associative strength could be accounted for by subjective ratings. Accuracy models were also compared with BIC scores. In this case, the subjective model (BIC = 4537.66) outperformed both the full model (BIC = 4547.15) and the objective-only model (BIC = 4555.36).

### 3.3. Discussion

Similarly to Experiment 1, these results suggest that the reported expectedness explains variance in participants’ perceptual decisions beyond that of objective model-based expectations – with both predictors explaining reaction time but the model including only ratings providing the best fit to accuracy.

## 4. Experiment 3

The aim of Experiment 3 was to replicate the basic effect of Experiment 2 and to probe a different subjective dimension by asking participants to rate surprise. Surprise may simply reflect the inverse scale of expectedness, but it is also possible that qualitatively different patterns could emerge because ‘surprise’ may represent a distinct construct that emerges only at the most extreme end of (low) subjective expectation.

### 4.1. Methods

#### 4.1.1. Participants

Thirty-five participants (*M* age = 31.27 years, *SD* = 6.72) completed the study and were included in the analysis. For Experiment 3, we pre-registered more stringent exclusion criteria: Participants whose accuracy at the orientation discrimination task was below 75% were excluded, as well as those at below 80% on catch trials. Three participants were excluded based on these criteria. This change in criteria had no effect on the pattern of results: Model results using the previous criteria (above chance) are reported in the supplementary materials. All participants reported normal or corrected-to-normal vision, normal hearing, and were recruited from Prolific. They gave their informed consent prior to participation and were paid an hourly wage of £9. The study complied with all relevant ethical regulations and was approved by the local Research Ethics Committee. The study design and analysis plan were pre-registered prior to data collection (OSF link: https://osf.io/y3xhc).

#### 4.1.2. Procedure

The experimental procedure was the same as that in Experiment 2 with some small changes (see Fig. 3B). The first change was the introduction of a fixed Gabor presentation time of 500 ms, as well as a fixed response period of 1500 ms after Gabor onset (1000 ms after offset) for participants to perform the orientation task. This was intended to standardise the duration of the visual stimulation of the Gabor and reduce the total duration of the experiment. The second set of changes related to the surprise scale. In Experiment 2, the expectation scale was a horizontal line with scale values increasing from 0 to 100 in increments of 1 from left to right. To decorrelate ratings from systematic motor responses, we now designed a surprise scale consisting of a spiral shape of increasing width. To rate their surprise, participants moved a red dot from the middle either to the thick end (for high surprise) or the narrow end (for low surprise) of the scale. The surprise scale ranged from 0 to 100 in increments of 10. The scale was rotated pseudo-randomly on each trial and flipped to the mirror image on half of trials.

#### 4.1.3. Analysis

RTs below 100 ms were excluded from the analysis along with trials in which the orientation discrimination question was answered incorrectly for the RT analyses. As the introduction of a response period meant the maximum reaction time was 1500 ms, there was no upper limit on reaction times in our analyses. In this experiment, we used the trial-wise prediction errors derived from the Rescorla-Wagner model as our measure of the objective task statistics. Surprise ratings were compared to objective prediction errors using linear mixed effects models, as in the previous experiments. Specifically, participants’ trial-by-trial ratings were predicted from the objective prediction error values using a linear mixed effects model. To investigate the influence of surprise and prediction error on behaviour, the same three models (objective, subjective, full) were used to predict RT and accuracy with participants’ trial-by-trial surprise ratings as the subjective predictor and RW model prediction error on each trial as the objective. In the accuracy models, the inclusion of trial number as a random effect again resulted in convergence issues – therefore, trial number was removed from the random effect structure of these models. It was retained as a fixed effect, as well as a fixed and random effect in reaction time models, as pre-registered.

### 4.2. Results

As in Experiment 2, we found that participants’ responses were significantly faster (*t*(34) = -5.749, *p* < .001) in the 75% condition (*M* = 564.43 ms, *SD* = 126.90) than the 25% condition (*M* = 619.78 ms, *SD* = 139.72). Likewise, discrimination accuracy was significantly higher (*t*(34) = 2.812, *p* = .008) for 75% trials (*M* = 95.39%, *SD* = 5.06) than 25% trials (*M* = 92.68%, *SD* = 5.98), suggesting that the design successfully induced learning about cue-orientation mappings.

Examining the coupling between subjective surprise and model-derived prediction error, revealed a significant positive association between prediction error and subjective surprise (*β* = 13.384, 95% CI [9.189, 17.580], *p* < .001), with higher model-derived prediction errors associated with higher subjective surprise ratings.

In reaction times, the subjective model revealed that as participants rated the stimulus as more surprising, they were slower to discriminate its orientation (*β* = 0.067, 95% CI [0.050, 0.084], *p* < .001). The objective model revealed that as the magnitude of objective prediction error increased, reaction times also slowed (*β* = 0.053, 95% CI [0.039, 0.067], *p* < .001). The full model showed contributions of both subjective surprise ratings (*β* = 0.045, 95% CI [0.023, 0.066], *p* < .001; Fig. 5C and D) and objective RW prediction error (*β* = 0.030, 95% CI [0.012, 0.047], *p* = .002; Fig. 5A and B) to RTs. As in the previous experiments, this significance of both predictors within the same model indicates that subjective and objective surprise explained independent variance in RTs. BIC scores marginally favoured the full model (BIC = 1506.42) over the subjective-only model (BIC = 1506.91). The negligible difference between the full and subjective model suggests they provide an equivalent fit to the data, but importantly both the full and subjective model outperformed the objective-only model (BIC = 1510.30).

**Figure 5:**
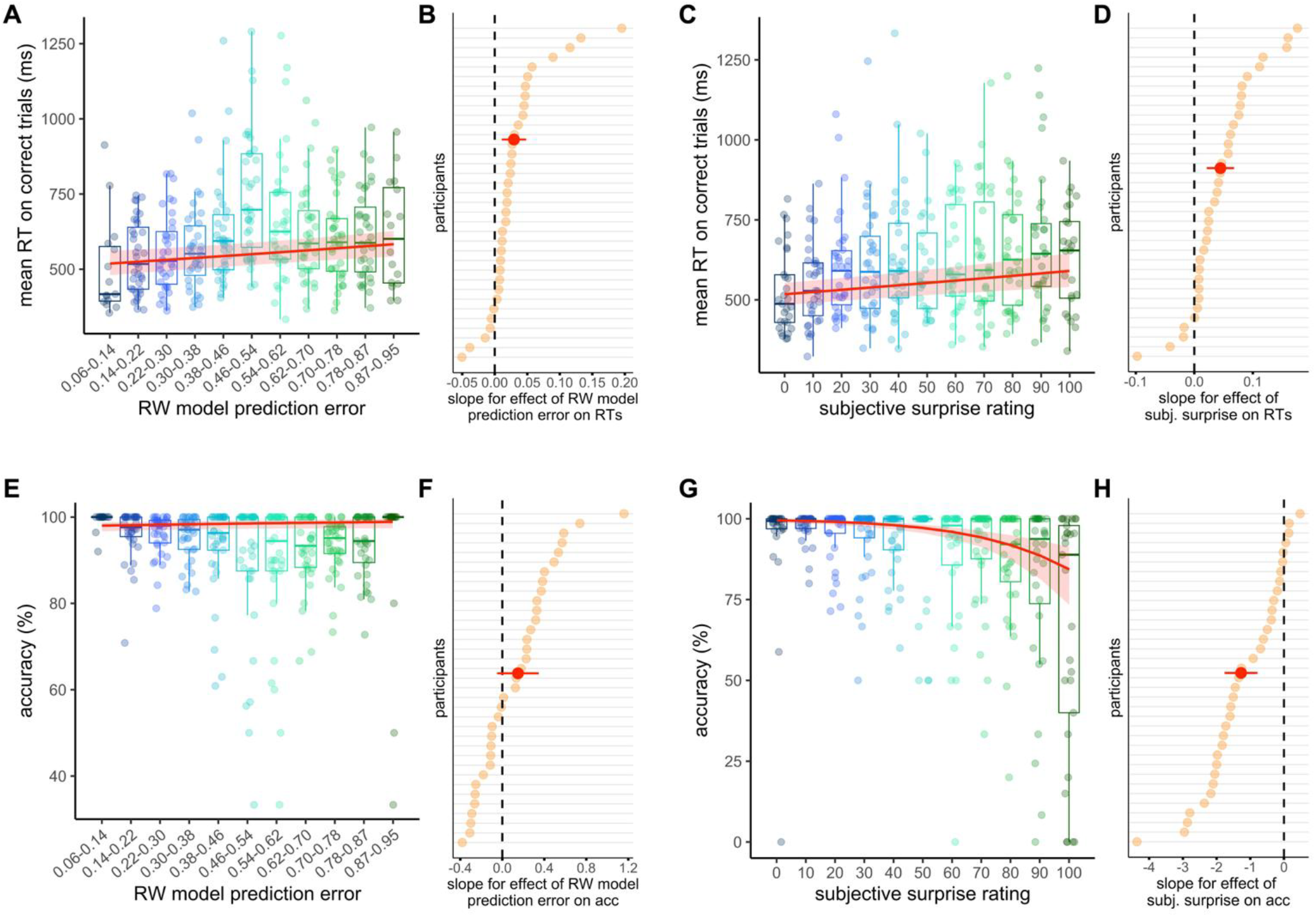
Experiment 3 results. **(A)** RTs as a function of Rescorla-Wagner prediction error, showing slower responses when the stimulus elicited a larger objective prediction error. **(B)** Participant-specific slopes for the effect of RW prediction error on RTs, with most participants showing a positive slope. **(C)** RTs as a function of subjective surprise ratings, indicating slower responses to outcomes rated as more surprising. **(D)** Participant-specific slopes for the effect of subjective surprise on RTs, with most participants exhibiting a positive relationship. **(E)** Accuracy as a function of RW prediction error. When included alongside subjective surprise ratings, objective RW prediction errors had no significant effect on accuracy. **(F)** Participant-specific slopes for the effect of objective RW prediction errors on accuracy. **(G)** Accuracy as a function of subjective surprise, showing decreasing accuracy as outcomes were rated as more surprising. **(H)** Participant-specific slopes for the effect of subjective surprise on accuracy, with the majority exhibiting a negative relationship. Plotting conventions are the same as for Fig. 4.

For accuracy, the subjective model showed that higher surprise ratings predicted lower accuracy (*β* = - 1.008, 95% CI [-1.312, -0.703], *p* < .001). The objective model replicated this effect for objective prediction error: larger model-derived prediction errors were associated with significantly reduced accuracy (*β* = - 0.248, 95% CI [-0.401, -0.095], *p* = .001). However, as in Experiment 2, in the full accuracy model objective values ceased to be significant (*β* = 0.149, 95% CI [-0.039, 0.338], *p* = .120; Fig. 5E and F), while subjective surprise still predicted accuracy (*β* = -1.273, 95% CI [-1.732, -0.813], *p* < .001; Fig. 5G and H). This pattern indicates that subjective surprise accounts for more variance in accuracy than objective prediction error. Accuracy model comparisons followed the general pattern of Experiments 1 and 2: The subjective model (BIC = 4550.23) best fit the data, compared to the objective (BIC = 4571.05) and the full (BIC = 4557.40) model.

### 4.3 Discussion

These findings replicate the patterns demonstrated in Experiment 2, but with ratings of surprise rather than expectations.

## 5. General Discussion

Across three studies, we asked about the role of subjective experiences of statistical structure in perceptual prediction. To this end, we investigated whether subjective reports of structure and objective statistics differentially explain reaction times and accuracy in perceptual decisions about a probabilistically cued stimulus. Experiment 1 assessed participants’ end-of-experiment estimates of cue-stimulus probabilities, while Experiments 2 and 3 examined how reports of statistical structure evolve during learning through trial-wise ratings of the expectedness (Experiment 2) and surprise (Experiment 3) associated with the cued stimulus. These subjective ratings were compared to the objective task structure – either ‘ground-truth’ probabilities (Experiment 1) or computationally-derived trial-wise estimates of associative strength (Experiment 2) and prediction error (Experiment 3). In all experiments, subjective ratings explained unique variance in perceptual decisions beyond that captured by objective structure, with the ratings explaining more unique variance in accuracy than objective structure itself.

Extensive research over the last three decades has shown that predictions exert a fundamental influence on a range of cognitive functions (Friston, 2018). However, previous studies have largely modelled statistical regularities and resulting expectations in terms of objective structure – e.g., binary categorisations of expected versus unexpected stimuli (e.g. Garlichs & Blank, 2024), or the probability of an event (Bestmann et al., 2008; Feuerriegel et al., 2021). By relying solely on objective structure, we risk missing meaningful variability in experience that could explain behaviour and neural responses but would remain hidden without explicit measurement. Specifically, subjective experience of statistical structure will presumably usually covary with the structure itself, and it did in the present studies. Therefore, while objective probabilities and learning models such as the RW can explain a great deal of variance in behaviour, modelling objective structure alone will render it impossible to disentangle the relative objective and subjective contributions. It is possible that modelling of subjective experience may also resolve some current debates in the literature, for example, concerning whether aberrant prediction generates the atypicalities characteristic of autism (Angeletos Chrysaitis & Seriès, 2023) or schizophrenia (Corlett et al., 2019), or whether it shapes early, or just later stages of, sensory processing (Gabhart et al., 2025; Thomas et al., 2024; Westerberg et al., 2025). For example, those studies finding effects may have generated differences in subjective experience, while nulls may be explained by objective, but not subjective, variation.

One particularly interesting possibility for the distinct behavioural variance explained is that subjective awareness of statistical structure, or error, drives qualitatively distinct processes that shape learning and perception. Many theories of awareness suggest that while objective ‘evidence’ throughout the brain can be continuous and graded, there is a qualitative difference between being consciously aware or not. For example, global workspace models propose an all-or-nothing ‘ignition’ for signals passing a threshold (Charles et al., 2013; Dehaene & Naccache, 2001), while likewise the activation of higher-order representations (Lau & Rosenthal, 2011), or recurrent dynamics in sensory structures (Lamme & Roelfsema, 2000), are generated in a discrete rather than graded fashion. If the function of experience is to enable model-based planning of action (Fleming & Michel, 2025), experience of surprise may be necessary for saccadic and manual exploration of the environment to resolve error – ‘exploration’ vs ‘exploitation’ modes (Cohen et al., 2007; Wyatt et al., 2024). Processes typically linked to objective prediction errors, such as neurochemical processes (e.g., Yu & Dayan, 2005) and the updating of an internal statistical model (Den Ouden et al., 2012), may also be, at least partially, related to the subjective experience that an event has violated a statistical regularity, rather than objective structure, per se.

Importantly, our argument does not imply that learnt contingencies are *only* able to influence behaviour and neural responses through subjective experiences of structure. While controversial, there is an extensive literature suggestive of implicit learning (Destrebecqz & Cleeremans, 2001; Turk-Browne et al., 2005, 2009), as well as the demonstration of neural and behavioural responses associated with both perceptual prediction (Batterink et al., 2024; Turk-Browne et al., 2010) and error (den Ouden et al., 2009) while participants cannot report the probabilistic relationships. However, just because learning and perceptual prediction perhaps can be realised implicitly, does not mean that it always, or even usually, is as such.

We were interested in how subjective ratings may dissociate from objective structure with an ultimate aim of interrogating the role of subjective experience. While variance in experience is likely to be explained well by self-report measures, report, and its dissociation from objective structure, will of course also reflect subpersonal and motoric or decisional processes (Tsuchiya et al., 2015). This is a problem that riddles any examination of awareness or experience, but self-report is still typically viewed as a good gold standard dependent variable by consciousness communities (Mudrik et al., 2025). It may be the best we currently have, especially given the wholesale differences in proposed neural signatures under distinct theories, and we believe is certainly a better reflection of experience than objective structure. Therefore, it was at minimum a good starting point for examinations. We also note that there may be other descriptors of objective structure than the particular ones we employed. Most notably, some may argue that models taking account of predictive hierarchical structure, and hence contextual variability, are often more fitting. We deemed the Rescorla-Wagner model the best choice in this setting given the structure did not vary across time, but extensions could compare results across different models. Importantly, the same conclusion was reached in Experiment 1 that determined objective probabilities by the global task structure, but even more importantly, there is no reason to think that subjective structure would perfectly track a different objective one – so our findings would remain relevant regardless of these relative comparisons.

In conclusion, we here demonstrate that subjective ratings of probability, expectedness, and surprise explain variance in perceptual prediction. This explanatory power was at least as strong, if not stronger, than the objective statistical structure, and thus opens a host of questions concerning their relative roles in predictive processes.

## Acknowledgements

This research was supported by a European Research Council (ERC) consolidator grant (101001592) under the European Union’s Horizon 2020 research and innovation programme awarded to CP.

## Data availability

Materials: All experimental materials are publicly available on the Open Science Framework (OSF). Data: All primary data are publicly available on OSF. Analysis scripts: All analysis scripts are publicly available on OSF. The OSF link for the entire project can be found at: https://osf.io/nte5u/files/osfstorage#.

## Notes

https://osf.io/nte5u/files/osfstorage#.

### Competing Interest Statement

The authors have declared no competing interest.

### Summary of Updates

The abstract and discussion have been slightly expanded.

