## Supplementary Materials for "Perceptual predictions track subjective, over objective, statistical structure"

**Model comparison: BIC vs AIC**

BIC scores strongly penalise for model complexity, favouring more parsimonious models. For completeness, we also examined AIC scores, which place greater emphasis on model fit relative to complexity. Results of BIC and AIC analyses are reported together below to aid comparison.

1. ***Experiment 1***

*1.1 Reaction time*

In the model comparison for Experiment 1 reaction time models, AIC and BIC scores were in agreement. Both showed that the full model – including both subjective and objective probability – provided the best fit (AIC = -9225.73, BIC = -9104.06), compared to the subjective-only (AIC = -9198.01, BIC = -9084.46) and objective-only (AIC = -9213, BIC = -9099.10) models.

*1.2 Accuracy*

In the model comparison for Experiment 1 accuracy models, AIC and BIC showed a slight divergence. BIC favoured the subjective model (BIC = 15763.50), outperforming the full (BIC = 15768.56) and objective (BIC = 15765.89) models. In contrast, AIC favoured the full model (AIC = 15653.65) over the subjective (AIC = 15656.80) and objective (AIC = 15659.19) models. Of importance however, given our conclusions, the objective only never won.

2. ***Experiment 2***

*2.1. Reaction time*

In the model comparison for Experiment 2 reaction time models, AIC and BIC scores were in agreement. Both metrics showed that the full model – including both subjective and model-derived expectedness – provided the best fit (AIC = 8017.16, BIC = 8128.76), followed by the subjective-only model (AIC = 8033.43, BIC = 8137.59), and then the objective-only model (AIC = 8048.518, BIC = 8152.68).

*2.2. Accuracy*

In the model comparison for Experiment 2 accuracy models, AIC and BIC scores also agreed. Both AIC and BIC showed that the **subjective model** provided the best fit to the accuracy data (**AIC = 4477.65, BIC = 4537.66**), outperforming both the objective model (**AIC = 4495.34, BIC = 4555.36**) and the full model including both predictors (**AIC = 4479.63, BIC = 4547.15**).

3. ***Experiment 3***

*3.1. Reaction time*

In the model comparison for Experiment 3 reaction time models, AIC and BIC scores again showed a consistent pattern. Both AIC and BIC indicated that the full model, which included both subjective and model-derived surprise, provided the best fit to the data (AIC = 1394.30, BIC = 1506.42). This model outperformed the subjective-only model (AIC = 1402.27, BIC = 1506.91) and the objective-only model (AIC = 1405.66, BIC = 1510.30).

*3.2. Accuracy*

In the model comparison for Experiment 3 accuracy models, AIC and BIC showed a slight divergence: BIC favoured the subjective model (BIC = 4550.23) over the full model (BIC = 4557.40), while again the objective model had the worst fit (BIC = 4571.05). In contrast, based on AIC the full model and the subjective model were equally well-fitting (full AIC = 4482.11; subjective AIC = 4482.48), while the objective model showed a poorer fit (AIC = 4503.29). Therefore, like with Experiment 1, most importantly for our conclusions and regardless of whether we used BIC or AIC, the objective model never won.

**Experiment 1 subjective surprise results**

Using participants’ surprise instead of probability ratings as the subjective predictor, produces similar results. When predicting reaction times, the subjective model revealed a positive effect of subjective surprise ratings (*β* = 0.027, 95% CI [0.013, 0.041]*,* *p* < .001), indicating that participants responded slower for stimuli they rated as more surprising. The objective model showed a negative effect of objective probability (*β* = -0.029, 95% CI [-0.036, -0.022]*,* p < .001), with faster RTs to objectively more probable outcomes. Importantly, the full model – containing both predictors – showed significant effects of both objective probability (*β* = -0.028, 95% CI [-0.035, -0.021]*,* p < .001; Fig. S1A and B) and subjective surprise ratings (*β* = 0.020, 95% CI [0.006, 0.034]*,* p = .007; Fig. S1C and D), suggesting that each explained independent variance in participants’ RTs. These results indicate that participants responded more slowly not only to outcomes that were objectively less probable, but also to those they rated as more surprising. However, model comparisons using BIC favoured the objective model (BIC = -9180.29) over the subjective (BIC = -9146.33) and full (BIC = -9177.47) models, whereas AIC favoured the full model (AIC = -9299.14) over the subjective (AIC = -9259.88) and objective (AIC = -9293.85) models.

When predicting accuracy, the subjective model revealed a negative effect of subjective surprise ratings (β = -0.201, 95% CI [-0.336, -0.066], p = .004), indicating that participants were less accurate for stimuli they judged as more surprising. The objective model showed a positive effect of objective probability (β = 0.150, 95% CI [0.062, 0.239], p < .001), suggesting higher accuracy for more objectively probable outcomes. In the full model, both subjective ratings (β = -0.158, 95% CI [-0.288, -0.028], p = .017; Fig. S1G and H) and objective probability (β = 0.125, 95% CI [0.039, 0.210], p = .004; Fig. S1E and F) remained significant, suggesting each explaining unique variance. Model comparison using BIC favoured the objective model (BIC = 15662.24) over the subjective (BIC = 15664.72) and full (BIC = 15667.07) models, whereas AIC favoured the full model (AIC = 15552.16) over the subjective (AIC = 15558.02) and objective (AIC = 15555.54) models.

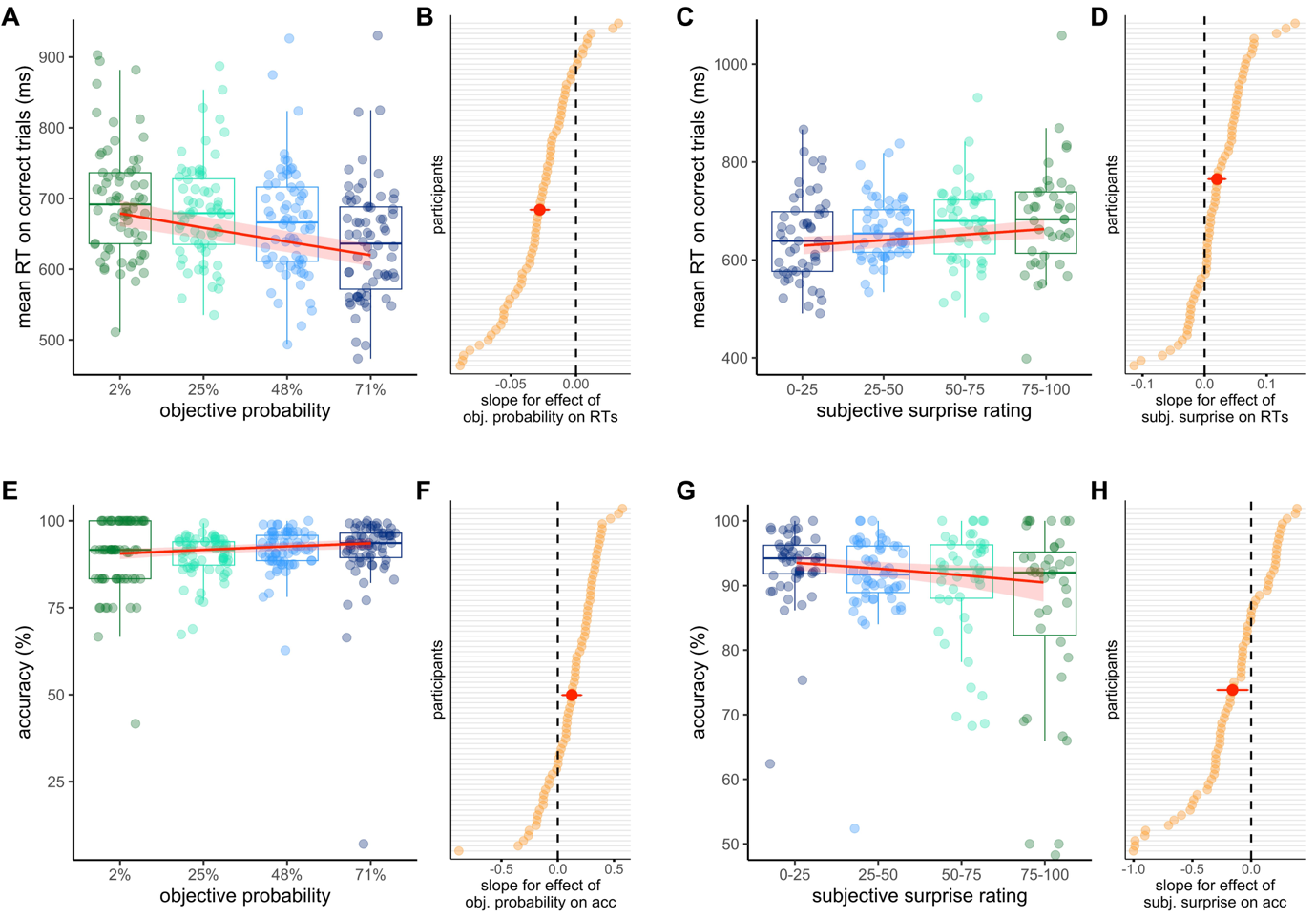

**Figure S1: Experiment 1 subjective surprise results. (A)** RTs as a function of objective probability, showing faster responses to outcomes that were objectively more probable. **(B)** Participant-specific slopes for the effect of objective probability on RTs, with most participants showing a negative slope. **(C)** RTs as a function of subjective surprise ratings, indicating faster responses to outcomes rated as less surprising. **(D)** Participant-specific slopes for the effect of subjective surprise on RTs, with most participants exhibiting a positive relationship. **(E)** Accuracy as a function of objective probability, showing fewer errors for outcomes that were objectively more probable. **(F)** Participant-specific slopes for the effect of objective probability on accuracy, with the majority showing a positive association. **(G)** Accuracy as a function of subjective surprise, showing fewer errors for outcomes rated as less surprising. **(H)** Participant-specific slopes for the effect of subjective surprise on accuracy, with the majority exhibiting a negative relationship. Plotting conventions are the same as for Fig. 2 in the paper.

**Experiment 3 results with different exclusion criteria**

To demonstrate that the more stringent exclusion criteria for Experiment 3 had no influence on the results, fixed effects estimates and model comparison scores using the same exclusion criteria as Experiment 1 and 2 (above chance accuracy and above 80% catch trial performance) (*N* = 38) are reported below.

In linear mixed-effects models predicting log-transformed reaction time, the subjective model showed that higher surprise rating was positively associated with reaction time (*β* = 0.059, 95% CI [0.041, 0.076], p < .001). In the objective model, prediction errors were also positively associated with reaction time (*β* = 0.045, 95% CI [0.031, 0.059], p < .001). In the full model including both predictors, both effects remained significant: surprise ratings (*β* = 0.039, 95% CI [0.019, 0.059], p < .001) and prediction errors (*β* = 0.028, 95% CI [0.013, 0.044], p = .001). This indicates that subjective surprise and prediction errors each independently contribute to predicting reaction time.

As in the reduced sample, model comparison using AIC and BIC confirmed that the full model (AIC = 2081.52, BIC = 2194.54) provided the best fit to the data, outperforming both the subjective model (AIC = 2090.43, BIC = 2195.91) and the objective model (AIC = 2091.24, BIC = 2196.72). These comparisons support the conclusion that both forms of surprise offer unique explanatory value.

In the accuracy models, trial number was removed as an additional fixed effect, as in the full sample this resulted in convergence issues (as was the case in Experiment 2). In the **subjective accuracy model,** surprise rating was a significant negative predictor of accuracy (*β* = -0.850, 95% CI [-1.069, -0.630], p < .001). In the **objective model,** prediction errors were also a significant negative predictor of accuracy (*β* = -0.189, 95% CI [-0.282, -0.095], p < .001). In the **full model** including both predictors, only subjective surprise remained significant (*β* = -0.984, 95% CI [-1.308, -0.660], p < .001), while prediction error was no longer a significant predictor (*β* = 0.067, 95% CI [-0.051, 0.185], p = .268).

Model comparison supported this pattern: the subjective model provided the best fit to the data, with the lowest AIC (7448.50) and BIC (7509.53), followed closely by the full model (AIC = 7449.25, BIC = 7517.91). The objective model was the least optimal fit (AIC = 7473.06, BIC = 7534.09).
